# Utilization of 5’-deoxy-nucleosides as Growth Substrates by Extraintestinal Pathogenic *E. coli* via the Dihydroxyacetone Phosphate Shunt

**DOI:** 10.1101/2023.08.10.552779

**Authors:** Katherine A. Huening, Joshua T. Groves, John A. Wildenthal, F. Robert Tabita, Justin A. North

## Abstract

All organisms utilize *S*-adenosyl-L-methionine (SAM) as a key co-substrate for methylation of biological molecules, synthesis of polyamines, and radical SAM reactions. When these processes occur, 5’-deoxy-nucleosides are formed as byproducts such as *S*-adenosyl-L-homocysteine (SAH), 5’-methylthioadenosine (MTA), and 5’-deoxyadenosine (5dAdo). One of the most prevalent pathways found in bacteria for the metabolism of MTA and 5dAdo is the DHAP shunt, which converts these compounds into dihydroxyacetone phosphate (DHAP) and 2-methylthioacetaldehyde or acetaldehyde, respectively. Previous work has shown that the DHAP shunt can enable methionine synthesis from MTA or serve as an MTA and 5dAdo detoxification pathway. Here we show that in Extraintestinal Pathogenic *E. coil* (ExPEC), the DHAP shunt serves none of these roles in any significant capacity, but rather physiologically functions as an assimilation pathway for use of MTA and 5dAdo as growth substrates. This is further supported by the observation that when MTA is the substrate for the ExPEC DHAP shunt, the sulfur components is not significantly recycled back to methionine, but rather accumulates as 2-methylthioethanol, which is slowly oxidized non-enzymatically under aerobic conditions. While the pathway is active both aerobically and anaerobically, it only supports aerobic ExPEC growth, suggesting that it primarily functions in oxygenic extraintestinal environments like blood and urine versus the predominantly anoxic gut. This reveals a heretofore overlooked role of the DHAP shunt in carbon assimilation and energy metabolism from ubiquitous SAM utilization byproducts and suggests a similar role may occur in other pathogenic and non-pathogenic bacteria with the DHAP shunt.

**Importance:** Acquisition and utilization of organic compounds that can serve as growth substrates is essential for pathogenic *E. coli* to survive and multiply. Ubiquitous enzymatic reactions involving *S*-adenosyl-L-methionine as a co-substrate result in the formation of the 5’-deoxy-nucleoside byproducts, 5’-methylthioadenosine and 5’-deoxyadenosine. All *E. coli* possess a conserved nucleosidase that cleaves these 5’-deoxy-nucleosides into 5-deoxy-pentose sugars for adenine salvage. The DHAP shunt pathway, which is found in ExPEC strains but neither in intestinal pathogenic nor commensal *E. coli,* enables utilization of 5’-deoxy-nucleosides and 5-deoxy-pentose sugars as growth substrates by ExPEC strains. This provides insight into the diversity of sugar compounds accessible by ExPEC strains in recalcitrant and nutrient-poor environments such as the urinary tract during infection. Furthermore, given the dihydroxyacetone phosphate shunt pathway appears to only support aerobic *E. coli* growth, this suggests an explanation as to why intestinal strains that primarily exist in anoxic environments lack this pathway.

## Introduction

Infection caused by *Escherichia coli* is a major global health concern with significant health and economic impacts [1–3] . Extraintestinal Pathogenic *E. coli* (ExPEC) strains can cause infection in niches outside of the gut [2, 4]. These include uropathogenic *E. coli* (UPEC), a major cause of urinary tract infections (UTIs), along with strains that cause blood, prostate, and mammary infections, as well as neonatal meningitis [1, 5]. While infection can be caused by invasion of ExPEC strains originating outside of the body, ExPEC strains also occupy niches within the intestinal microbiome and when presented with the right conditions can cause opportunistic infections [2, 5, 6].

The ExPEC lineage initially diverged from commensal *E. coli* over 32 million generations ago [7], and ExPEC strains today appear diverse, harboring numerous horizontally acquired pathogenicity and fitness genes located within genomic islands [8, 9]. While numerous genomic island virulence and fitness genes have been identified that are involved in adhesion, iron uptake, and toxin production [3, 4, 9], many others remain unknown in their function or physiological role [10–12]. Previously using comparative genomics, we observed that nearly 50% of ExPEC strains, including the globally disseminated ST131 lineage, possess a conserved gene cluster known as the dihydroxyacetone phosphate (DHAP) shunt that is also found in a variety of environmental and pathogenic bacteria for the salvage of 5’-deoxy-nucleosides and 5-deoxy-pentose sugars (Fig. 1B-C) [13]. The *E. coli* variation of the DHAP shunt is composed of a nucleosidase (Pfs, known as MtnN in other organisms), a kinase (MtnK), an isomerase (MtnA), and an aldolase (Ald2) that function to convert 5’-methylthioadenosine (MTA) and 5’-deoxyadenosine (5dAdo) into adenine, DHAP, and an aldehyde (Fig. 1B-C). The other known DHAP shunt variation, which does not appear in *E. coli*, employs a bifunctional nucleosidase/phosphorylase (MtnP) that replaces the nucleosidase and kinase [13–15]. All *E. coli* possess the Pfs nucleosidase for purine salvage and the disposal of MTA and 5dAdo to prevent inhibitory buildup (Fig. 1B) [16–18]. In ExPEC strains, the remainder of the DHAP shunt is located as a single conserved gene cluster at the end of the tRNA-*leuX* genomic island [19]. To date, the DHAP shunt has not been identifiable in intestinal pathogenic or commensal *E. coli* strains [13]. For environmental freshwater and soil bacteria like *Rhodospirillum rubrum* and *Rhodopseudomonas palustris*, the DHAP shunt was shown to function physiologically as a sulfur salvage pathway. In these organisms, MTA was metabolized to 2-methylthioacetaldehyde for the synthesis of the essential amino acid methionine [14, 20–22] (Suppl. Fig. S1). In the insect pathogen, *Bacillus thuringiensis*, which also possesses the Universal Methionine Salvage Pathway for MTA recycling to methionine (Suppl Fig. S1B), the DHAP shunt was observed to serve as a detoxification and disposal pathway for 5dAdo [15]. However, the physiological role of the DHAP shunt in ExPEC has remained largely unknown.

**Fig. 1.**
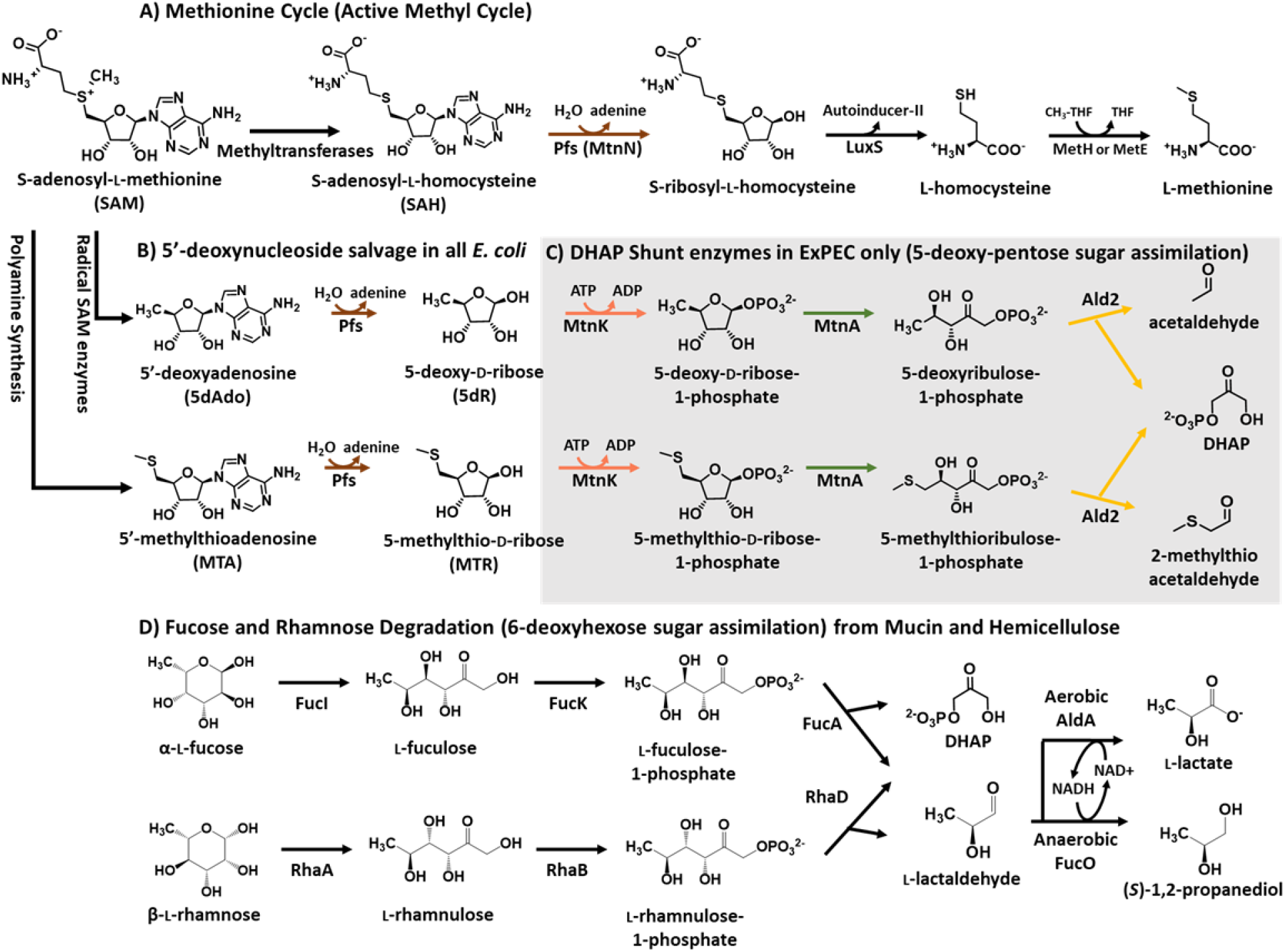
Salvage of SAM utilization byproducts, 5-deoxy-pentose sugars, and 6-deoxy-hexose sugars in *E. coli*. **A)** *S*-adenosyl-L-homocysteine (SAH), produced by methyltransferases is recycled by the Methionine Cycle (a.k.a. Active Methyl Cycle) [64, 65]. **B-C)** The *E. coli* variation of the DHAP shunt. B) For adenine salvage and detoxification of 5’-methylthioadenosine (MTA) and 5’-deoxyadenosine (5dAdo), all *E. coli* possess the multifunctional SAH/MTA/5dAdo nucleosidase (Pfs; a.k.a. MtnN) [23]. C) ExPEC strains possess the Pfs nucleosides (B) and remainder of the DHAP shunt genes which together compose a dual-purpose pathway for conversion of 5’-deoxy-nucleosides in the form of MTA and 5dAdo or corresponding 5-deoxy-pentoses in the form of 5-deoxy-D-ribose and 5-methylthioribose, respectively, into the central carbon metabolite dihydroxyacetone phosphate (DHAP) and acetaldehyde or (2-methylthio)acetaldehyde [13]. **D)** *E. coli* analogously metabolizes 6-deoxy-hexose sugars in the form of L-fucose and L-rhamnose to DHAP and L-lactaldehyde. During anaerobic growth L-lactaldehyde is primarily reduced to (*S*)-1,2-propanediol as a terminal product, whereas during aerobic growth it is primarily oxidized to L-lactate for carbon assimilation as pyruvate. **Enzymes**: **Pfs** (MtnN), SAH/MTA/5dAdo nucleosidase, E.C. 3.2.2.9; **LuxS**, *S*-ribosyl-L-homocysteine lyase, E.C. 4.4.1.21; **MetH**, methionine synthase, E.C. 2.1.1.13; **MetE**, methionine synthase, E.C. 2.1.1.4; **MtnK**, 5-methylthioribose/5-deoxyribose kinase, E.C. 2.7.1.100; **MtnA**, 5-methylthioribose-1-phosphate/5-deoxyribose-1-phosphate isomerase, E.C. 5.3.1.23; **Ald2**, 5-methylthioribulose-1-phosphate/5-deoxyribulose-1-phosphate aldolase, E.C. 4.1.2.62; **FucI**, L-fucose isomerase, E.C. 5.3.1.25; **FucK**, L-fuculose kinase, E.C. 2.7.1.51; **FucA**, L-fuculose-1-phosphate aldolase, E.C. 4.1.2.17; **RhaA**, L-rhamnose isomerase, E.C. 5.3.1.14; **RhaK**, L-rhamnulose kinase, E.C. 2.7.1.5; **RhaD**, L-rhamnulose-1-phosphate aldolase, E.C. 4.1.2.19; **FucO**, (*S*)-1,2-propanediol oxidoreductase, E.C. 1.1.1.77; **AldA**, L-lactaldehyde dehydrogenase, E.C. 1.2.1.22.

MTA, 5dAdo, and *S*-adenosyl-L-homocysteine (SAH) are metabolic byproducts formed by all organisms via enzymatic reactions using *S*-adenosyl-L-methionine (SAM; Fig. 1). Metabolites SAH, MTA, and 5dAdo are competitive inhibitors of SAM-utilizing enzymes and concentrations above 1 mM are inhibitory to *E. coli* growth [17, 18, 23]. Therefore, cells must either convert them to non-inhibitory compounds and/or excrete them outside of the cell [16]. In commensal *E. coli*, SAH is initially cleaved by the MTA/5dAdo/SAH nucleosidase (Pfs) to produce adenine and *S*-ribosylhomocysteine, which can continue in the methionine cycle to regenerate methionine and SAM (Fig. 1A) [23]. In contrast, commensal *E. coli* only partially recycles MTA and 5dAdo. Analogous to SAH, adenine is cleaved from MTA and 5dAdo by Pfs, but the resulting 5-deoxy-pentoses, 5-methylthio-D-ribose (MTR) and 5-deoxy-D-ribose (5dR), respectively, are excreted from the cell (Fig. 1B) [16, 17, 24].

Previously we reported that the DHAP shunt was active in *E. coli* ATCC 25922, an attenuated ExPEC strain originally isolated from a clinical patient in Seattle, Washington [25, 26], for both MTA and 5dAdo metabolism. However, the physiological role of the DHAP shunt was unknown and it was unclear why this pathway was present only in ExPEC strains. Here we report that *E. coli* ATCC 25922 uses the DHAP shunt as a means of carbon assimilation from externally acquired 5’-deoxy-nucleosides and 5-deoxy-pentose sugars for growth, analogous to 6-deoxy-hexose sugar metabolism by the fucose and rhamnose catabolic pathways of *E. coli* (Fig.1B-D). Furthermore, transfer of the DHAP shunt gene cluster to commensal strain K12 is sufficient to enable aerobic growth with a 5-deoxy-pentose sugar like 5dR. This distinguishes ExPEC’s use of this pathway from other environmental organisms previously described, which employ the DHAP shunt for sulfur salvage (Suppl. Fig. S1) [13, 14, 20] or 5-deoxy-pentose detoxification [15]. Finally, the DHAP shunt only supports aerobic *E. coli* growth from 5’-deoxy-nucleosides and 5-deoxy-pentoses, providing insights as to why intestinal pathogenic and commensal *E. coli* strains inhabiting the primarily anoxic human gut do not possess this pathway.

## Materials and Methods

### Fine Chemicals

All chemicals unless otherwise stated were from Millipore-Sigma. All gases for growth studies were of > 99.99% purity from Praxair. All isotopically labeled standards were synthesized exactly as previously described [13, 14].

### Strains and growth conditions

#### Bacterial Strains

*E. coli* clinical isolate ATCC 25922 (Seattle 1946) [25, 26] was used as an attenuated ExPEC strain containing the DHAP shunt genes. BW25113, parent strain of the Keio Collection which is derived from K-12 *E. coli* [27], was used as a model commensal strain. Deletion of gene cluster *mtnK-mtnA-ald2* in ATCC 25922 (strain ΔK2; Δ*mtnK* Δ*mtnA* Δ*ald2*) and the *pfs* gene were performed using the λ-red system [28] as previously described [13]. Primers used for PCR amplification from pKD4 for each gene deletion are listed in Suppl. Table S1. *E. coli* Stellar^TM^ (Takara Biosciences) was used for cloning and plasmid storage. All transformations were performed by electroporation.

#### General Growth

Cells were either grown in Lysogeny Broth (LB) or in modified M9 media as previously described [13], supplemented with 1 mg/L nicotinic acid (ATCC 25922 is an auxotroph), 40 mM sodium nitrate (as an electron acceptor for anaerobic respiration), 1 mM ammonium sulfate unless otherwise indicated, and either glucose, 5dR, MTA, or 5dAdo at indicated concentrations. Aerobic and anaerobic cultures were incubated at 37 °C with shaking at 250 rpm. For liquid growth experiments, ATCC 25922 was first grown overnight aerobically in liquid LB cultures, then washed three times in carbon-free or sulfur-free M9 media prior to inoculation in M9 media to an initial optical density at 600 nm (OD_600nm_) of ∼0.05. Anaerobic manipulations were performed in an anaerobic chamber with 5%/95% H_2_/N_2_ atmosphere (Coy Laboratories).

#### Growth Inhibition Studies

*E. coli* strains were grown aerobically in M9 with 25 mM glucose to mid-exponential phase, washed once with M9 glucose media and used to inoculate 2 ml M9 glucose media supplemented with MTA, 5dAdo, or 5dR to an initial OD_600nm_ of ∼ 0.2. Cells were grown aerobically for 18 hours to allow each culture to reach its final maximum optical density. LD_50_ values were determined by weighted nonlinear parametric fit (MatLab, Mathworks) to the Hill equation using the average and standard deviation of final culture density versus inhibitor concentration for n=3 independent experiments.

#### Oxygen Limitation

Anaerobic cultures tubes containing M9 media with either 25 mM glucose, 5 mM 5dR, or no carbon source in were inoculated, stoppered and seald, and fitted with 18 Ga needles as an inlet and outlet. For 15 minutes, the culture was sparged with an atmosphere of defined oxygen concentration made by mixing purified air together with N_2_ via a gas mixer (Matheson #602; 0 PSIG) at a rate of 100-200 μl/s and 1 atm total pressure. Cultures were incubated at 37 °C in a shaking water bath at 120 rpm and the headspace was purged with the same gas mixture. The flow rate for each gas (air and nitrogen) and coordinately the O_2_ partial pressure was calculated using the manufacturer’s gas flow rate calibration charts (Matheson #602; 0 PSIG).

#### Serial Plating and Growth

Solid agar M9 plates for serial dilution growth studies were made with 1 mM glucose, MTA, 5dAdo, or 5dR; 7.5 g/L Noble agar (Affymetrix) and molecular biology grade water (Sigma). Cells were initially grown aerobically in 5 ml M9 media supplemented with 5 mM glucose to mid-exponential phase, washed 4 times with carbon-free M9 media, serially diluted onto the plates, and incubated at 37 °C for 2-3 days.

#### Plasmid Construction

An *E. coli* complementation vector, pTETTET, with a tetracycline-inducible promoter was constructed by replacing the *lac* promoter of pBBRsm2-MCS6 [29] with the dual *tet* promoter plus *tetR* and the 70 bp region immediately downstream of *tetR* from the transposon Tn*10* [30, 31] as fully detailed in the SI Methods. The *mtnK-mtnA-ald2* gene cluster was then cloned into pTETTET by PCR amplification using primers K2-NdeI-F and K2-SacI-R (Suppl. Table S1) that introduced flanking NdeI and SacI sites. This resulted in plasmid pK2 for tetracycline-inducible expression of DHAP shunt genes in *E. coli*.

#### Targeted analysis of DHAP shunt metabolites

HPLC analysis of cellular SAH, MTA, and 5dAdo was performed as previously described [13]. Targeted metabolomics of aerobic *E. coli* strains fed with [^14^C-methyl]-5’-methylthioadenosine or [5’-^3^H]-5’-deoxyadenosine was also performed as previously described [13] as fully detailed in the SI Methods .

#### Untargeted Metabolomics

Untargeted metabolomics were performed at the Ohio State University Campus Chemical Instrument Center. *E. coli* ATCC 25922 was grown in M9 media with 25 mM glucose, 1 mM sulfate, and supplemented with or without 1 mM MTA. Cells were grown aerobically at 37 °C with shaking at 250 rpm to mid-exponential phase and harvested by centrifugation. The culture media was retained and cells were extracted by an equal volume of 50% acetonitrile in water. The combined media and extracted metabolites were lyophilized and resuspended in 50% acetonitrile in water for LC-MS/MS analysis. Metabolites were resolved using an Agilent ZORBAX SB-Aq C18 reverse phase column (3.0 x 150 mm, 3.5 μm resin) on a Dionex/Thermo UltiMate 3000 HPLC System. Metabolites were eluted on a linear gradient of 5% to 20% acetonitrile in water plus 0.1% formic acid. Full mass and MS/MS were acquired by ESI in the negative mode on a Thermo QE M UPLC MS/MS System.

#### Cell Yield Measurements

*E. coli* ATCC 25922 was grown aerobically in M9 media with 5 mM of each growth substrate. Cells were harvested by centrifugation, spent media was retained, and the cell pellets were lyophilized to dryness to quantify total cell mass. The initial and final concentration of each substrate before and after growth, respectively, was quantified by C18 reverse phase HPLC using phenylhydrazine derivatization. To each 500 μl sample of spent media, 10 μl of fresh 50 mg/ml phenylhydrazine in 10% glacial acetic acid was added and incubated at 100 °C for 1 hr with occasional vortexing. Metabolites were resolved on a gradient of 17.5% - 50% acetonitrile in water with 20 mM ammonium acetate over 20 minutes with UV detection at 340 nm. Metabolite identities and concentrations were determined based on standard calibration curves of known metabolite standards. Cell yield (Y_max_) was calculated as the grams dry cell weight per gram substrate consumed.

## Results

### The DHAP shunt is not a significant means of sulfur acquisition in E. coli and leads to accumulation of 2-methylthioethanol and 2-methylsulfinylethanol

Seminal studies on the physiological role of the DHAP shunt in environmental photosynthetic bacteria like *Rhodospirillum rubrum* and *Rhodopseudomonas palustris* revealed that this pathway enabled sulfur salvage from MTA to maintain cellular methionine and cysteine pools [13, 14, 20, 22] (Fig. 1). This occurred through the further metabolism of the terminal DHAP shunt metabolite, 2-methylthioacetaldehyde, into 2-methylthioethanol and subsequent cleavage into methanethiol for methionine synthesis (Suppl. Fig. S1). As a result, *R. rubrum* and *R. palustris* could acquire sulfur and grow using MTA as a sole sulfur source via the DHAP shunt [20, 22]. For ExPEC strain ATCC 25922, initial work showed that the DHAP shunt was active for the metabolism of MTA to 2-methylthioethanol (Fig. 1C) [13]. However, it remained unclear as to whether ExPEC could salvage sulfur from MTA for methionine synthesis under aerobic or anaerobic conditions given that ExPEC strains are missing one or more gene homologs for any one of the known methionine salvage pathways present in other organisms (Suppl. Fig. S1) [13, 14, 20–22, 32–34].

To determine if the ExPEC DHAP was involved in salvage sulfur from SAM utilization byproducts, we performed growth studies with MTA or 2-methylthioethanol as a sole sulfur source (Fig. 2A). While ATCC 25922 was able to grow using sulfate or methionine as the sole sulfur source, it was completely incapable of growth with MTA or 2-methylthioethanol. This suggests that even if the DHAP shunt enables some sulfur salvage in the ATCC 25922 ExPEC strain, it is insufficient to support growth unlike in other organisms where the DHAP shunt has been established to function in distinct sulfur salvage pathways for MTA (Suppl. Fig. S1D) [14, 20, 22]. Similarly, we assessed whether the DHAP shunt was involved in recycling endogenously produced SAM utilization byproducts. Total amounts of MTA, 5dADO, and SAH produced during growth were determined by deletion of the *pfs* nucleosidase gene, which prevents their metabolism [17, 35]. ExPEC strain ATCC 25922 and commensal K12 produced similar amounts of SAM utilization byproducts (Suppl. Fig. S3A-B and SI results), and MTA levels (6-10 μmol/L/OD_600_) were similar to those previously reported in gut and commensal *E. coli* strains due to polyamine synthesis [36]. This indicates that ExPEC strains do not produce substantially more SAM utilization byproducts than their intestinal counterparts. Coordinately, when the ExPEC DHAP shunt genes were deleted (ATCC 26922 strain ΔK2; Δ*mtnK*/Δ*mtnA*/Δ*ald2*), there was no defect in growth when sulfate was limiting compared to the wild-type strain (Suppl. Fig. S3C-D). Together, this indicates that the DHAP shunt is not significantly involved in salvage of endogenously produced MTA, which is in stark contrast to *R. rubrum* and *R. palustris* [13, 14].

**Fig. 2.**
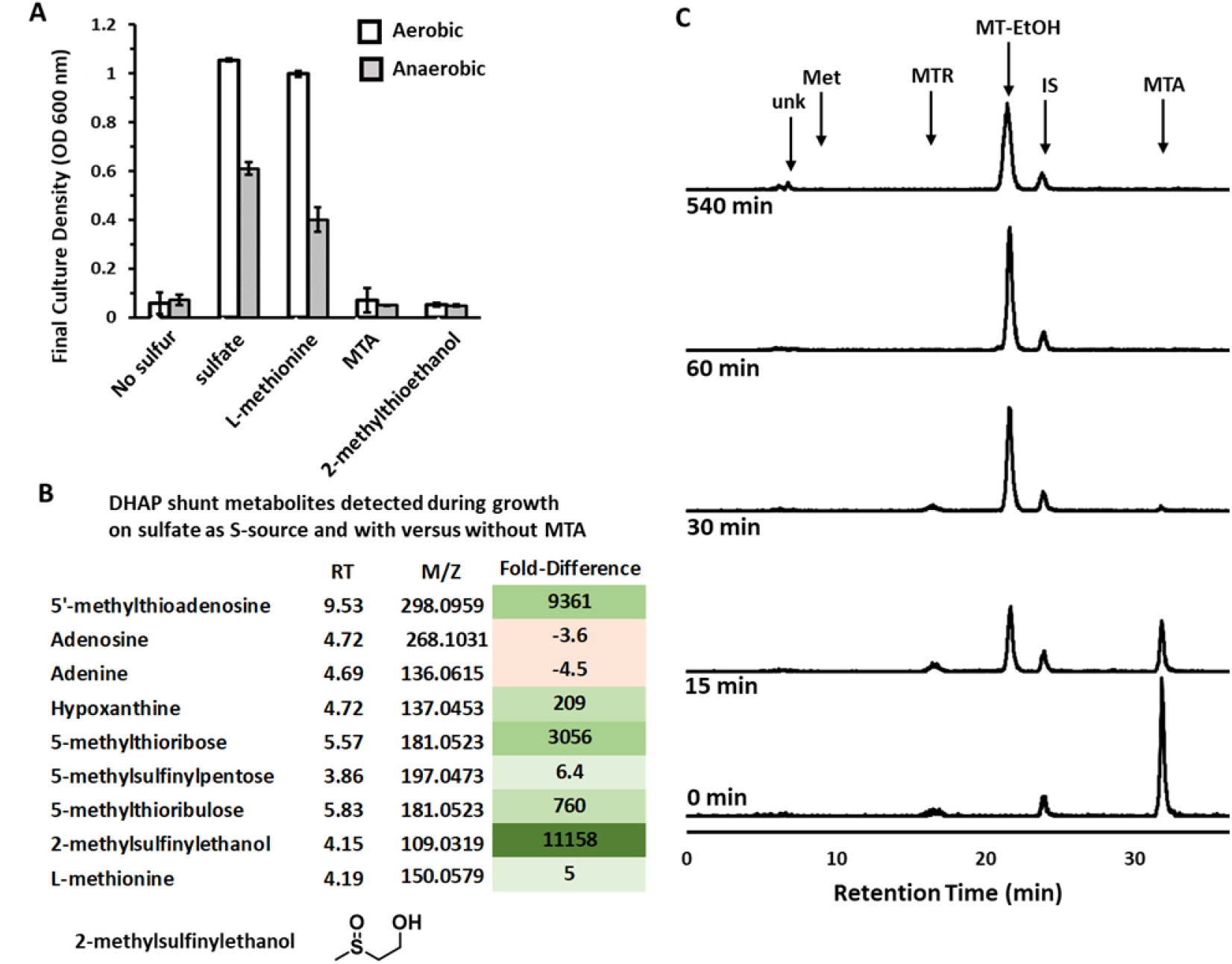
Sulfur from MTA cannot be salvaged by *E. coli* for growth. **A)** ATCC 25922 maximum growth achieved after 24 hours measured by optical density at 600 nm when cultured with 1 mM of the indicated sulfur compound as the sole sulfur source. No further growth was observed after 24 hours. Average and standard deviation error bars are for n=3 independent replicates. **B)** Fold-difference in the abundance of DHAP shunt-associated metabolites when ATCC 25922 was grown aerobically in the presence of 1 mM sulfate and 1 mM MTA versus grown aerobically in the presence of 1 mM sulfate only. Metabolites were resolved by LC-MS/MS from three independent biological replicates for each growth condition. Values are the average for the three replicates and significance of fold-change was analyzed by ANOVA with P < 0.05. **C)** Reverse Phase HPLC quantification of *E. coli* ATCC 25922 metabolites when fed aerobically with [^14^C-methyl]-5’-methylthioadenosine. Unk – unknown (identified as 2-methylsulfinylethanol by LC-MS/MS, m/z = 109.0319), R_T_ = 6.7 min; Met – methionine, R_T_ = 8.0 min; MTR – methylthioribose, R_T_ = 16.7 min; MT-EtOH – 2-methylthioethanol, R_T_ = 21.7 min; IS - internal standard, R_T_ = 24.0 min; MTA – 5’-methylthioadenosine, R_T_ = 31.8 min.

To further probe the metabolism of MTA by ExPEC strain ATCC 25922, particularly for methionine salvage and production of additional compounds during aerobic growth, we performed untargeted LC-MS/MS metabolomics of ATCC 25922 grown using sulfate supplemented with or without 1 mM MTA, and targeted metabolomics of cells fed with isotopically labeled MTA. When grown in the presence of or fed with MTA, there was a large increase in the abundance of DHAP shunt metabolites (Fig. 2B-C; MTA, MTR, 5-methylthioribulose). Concomitantly, hypoxanthine, which is the deaminated form of adenine, was the only purine that markedly increased in abundance, indicating that excess adenine cleaved from the MTA accumulates as hypoxanthine as previously observed in soil and freshwater bacteria [21]. Methionine, however, only increased in abundance by ∼ 5-fold when cells were grown in the presence of MTA (Fig. 2B), and no labeled methionine was detected after feeding with MTA (Fig. 2C; limit of detection of 1 nCi = 15 pmoles). This indicates that if methionine is salvaged from MTA, it is at low concentrations (< 150 nM). Rather the change in methionine abundance observed during growth in the presence of MTA is likely due to altered rates of *de novo* methionine synthesis and utilization given MTA is a competitive inhibitor of SAM-dependent enzymes and a regulator of cell metabolism [21, 23, 37]. Thus, the growth and metabolite analyses and the lack of identifiable gene homologs for any of the known methionine salvage pathways for MTA based on the DHAP shunt (Suppl. Fig. S1D) [13] all support the conclusion that the DHAP shunt is not a significant means of sulfur salvage in ExPEC, and any salvage is likely due to unknown, promiscuous processes.

Ultimately for sulfur metabolism from MTA by the DHAP shunt, both the untargeted and targeted metabolomics reveal that 2-methylthioethanol is the primary terminal product (Fig. 2B-C). After 30 minutes post feeding with MTA, 100% of the MTA was converted to 2-methylthioethanol, and even after 8 hours post feeding 2-methylthioethanol and a new species (R_T_ = 6.7 min) were observed. The slow conversion of 2-methylthioethanol (R_T_ =21 min) to the new species was found to be non-biological in nature by incubating labeled [2-methyl-^14^C]-methylthioethanol in media aerobically over the same time-course as cell feeding and resolving by reverse phase HPLC (Suppl. Fig. S4B). Coordinately, the untargeted metabolomics identified a compound with mass corresponding to 2-methylsulfinylethanol (m/z = 108.0246 Da, 108.0245 expected, error = 1 ppm) that highly increased in abundance when cells were grown in the presence of MTA (Fig. 2B). The reason 2-methylthioethanol was not observed by untargeted metabolomics is because of removal during the metabolite extraction and lyophilization process for LC-MS/MS (Fig. S4A). We verified that the extraction process does not cause oxidation of 2-methylthioethanol (Fig. S4A), confirming that the formation and observation of 2-methylsulfinylethanol is due to the slow non-enzymatic oxidation of 2-methylthioethanol after being formed by *E. coli* under oxic conditions (Fig. S4B). This previously unobserved non-enzymatic oxidation of 2-methylthioethanol is specific to aerobic conditions, as there is none observed during anaerobic growth [14, 22]. Ultimately, the 2-methylthioethanol produced by *E. coli* via the DHAP serves primarily as a terminal product that can slowly oxidize during aerobic growth conditions to 2-methylsulfinlyethanol.

### The ExPEC DHAP shunt is not a SAM utilization byproduct detoxification pathway

All *E. coli* possess the conserved Pfs enzyme, a multifunctional SAH, MTA, and 5dAdo nucleosidase [23]. This functions to cleave the SAM utilization byproducts into adenine and the corresponding ribosyl species (Fig. 1A-B). In the absence of a functional *pfs* gene, accumulation of SAH, MTA, and 5dAdo competitively inhibits the activity of SAM-dependent enzymes, leading to slower cell growth [17, 38, 39]. Historically for *E. coli*, the cleavage of SAH, MTA, and 5dAdo by the Pfs nucleosidase has been considered to be the sole and sufficient means for eliminating these inhibitory compounds [23]. *S*-ribosyl-L-homocysteine produced from SAH by Pfs is recycled into autoinducer-2 and methionine by the active methyl cycle (Fig. 1A). Conversely, MTR and 5dR generated by the cleavage of MTA and 5dAdo, respectively, are excreted as terminal byproducts (Fig. 1B) [23]. Recently, a possibly long-overlooked inhibitory role of MTR and 5dR was observed [15]. In the insect pathogen *Bacillus thuringiensis*, the DHAP shunt kinase, isomerase, and aldolase were required for maximal growth in glucose media if exogenously supplied 5dR was present. At concentrations of 1 mM 5dR, a ∼25% reduction in growth rate was observed for DHAP shunt deletion strains compared to the wild type [15]. This led to the hypothesis that the DHAP shunt’s role is to detoxify internally produced or exogenously present MTR and 5dR, providing a potential growth advantage for ExPEC strains over their commensal counterparts.

To assess the inhibitory effects of MTA, 5dAdo, MTR, and 5dR on ExPEC and commensal *E. coli* growth, and the physiological role of the conserved Pfs nucleosidase versus the ExPEC DHAP shunt, we performed growth inhibition studies (Fig. 3A-C). In both the wild type ATCC 25922 ExPEC and K12 commensal strains, MTA and 5dAdo were inhibitory to growth at concentrations above 2 mM. When the *pfs* gene was inactivated in either strain, sensitivity to MTA and 5dAdo increased 2- and 5-fold, respectively (Fig. 3D). In stark contrast, when the ExPEC DHAP shunt *mtnK, mtnA,* and *ald2* were deleted (Fig. 3A-B; strain ΔK2), there was no statistically significant change in sensitivity to MTA and 5dAdo, indicating the DHAP shunt MtnK, MtnA, and Ald2 did not aid in managing the inhibitory effects of MTA and 5dAdo, and by extension MTR and 5dR. This establishes the essentiality of the Pfs nucleosidase in ExPEC strains is to prevent the inhibitory buildup of SAM utilization byproducts as observed in commensal strains [17, 38]. We also directly quantified if 5dR was inhibitory. Even at a concentration of 4 mM supplied 5dR, there was no significant inhibitory effect on the growth of either ATCC 25922 or K12 (Fig. 3C). Furthermore, deletion of the ExPEC DHAP shunt by virtue of inactivating *mtnK, mtnA,* and *ald2* had no effect on ATCC 25922 sensitivity to 5dR (Fig. 3C). Thus, 5dR is not inhibitory in the millimolar range in which it can accumulate due to microbial metabolism [16].

**Fig. 3.**
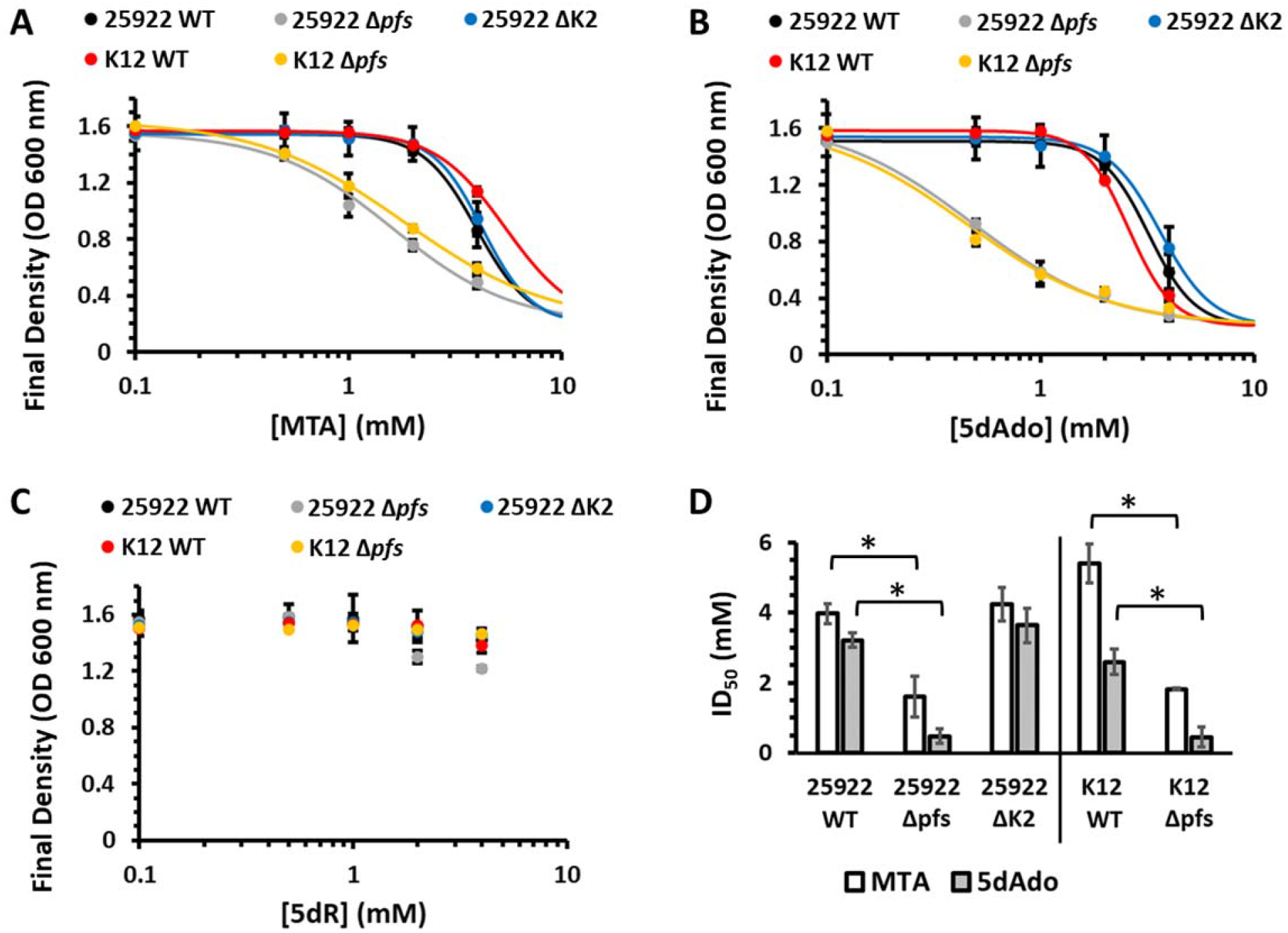
Prevention of growth inhibition by SAM utilization byproducts requires Pfs but not the DHAP shunt. Final culture density after 18 hours of the ATCC 25922 wild type, Δ*pfs*, and ΔK2 (Δ*mtnK* Δ*mtnA* Δ*ald2*) strains and the K12 BW25113 wild type and Δ*pfs* deletion strains grown aerobically with glucose and in the presence of the indicated concentration of either **A)** 5’-methylthioadenosine (MTA), **B)** 5’-deoxyadenosine (5dAdo), or **C)** 5-deoxy-D-ribose (5dR). Averages and standard deviation error bars are for n=3 independent replicates. Curves are the non-linear least squares weighted fit to the Hill equation. **D)** ID_50_ for MTA and 5dAdo in ExPEC ATCC 25922 and commensal K12 BW25113 strains. ID_50_ values are from the fit curves in A-B to the Hill Equation and the error bars are the parameter 95% confidence interval of the weighted fit. * Statistically significant difference, P < 0.05.

### The ExPEC DHAP shunt metabolizes SAM utilization byproducts for use as cellular growth substrates

As with commensal *E. coli,* many bacteria cannot utilize MTA and 5dAdo produced by SAM-dependent reactions beyond adenine salvage via the Pfs (MtnN) nucleosidase, resulting in the accumulation of the 5-deoxy-pentoses, MTR and 5dR (Fig. 1) [16, 17, 23, 24].

Furthermore, eukaryotic cells produce a diversity of 5’-deoxy-nucleosides as a byproduct of metabolism. These include MTA, 5’-methylthioinosine (MTI), 5dAdo, 5’-deoxyinosine (5dI), and 5’-deoxyxanthosine (5dX). In humans, these modified nucleosides each accumulate in the urine at concentrations of 0.1-10 μmol/mmol creatine [40–43]. Notably for *E. coli*, the Pfs nucleosidase is active with each of these 5’-deoxy-nucleosides resulting in the formation of MTR or 5dR [40]. Thus, the DHAP shunt is poised to be able to assimilate carbon from a wide number of 5’-deoxy-nucleosides and the 5-deoxy-pentoses, MTR and 5dR.

Initially, we assessed the ability of ExPEC strain ATCC 25922 to grow using 5dR. The strain grew aerobically with 5dR as a sole carbon source (Fig. 4A), but it could not grow anaerobically with 5dR either under fermentative conditions (not shown) or anaerobic respiratory conditions with nitrate (Suppl. Fig. S5). When the DHAP shunt was deleted by the inactivation of the *mtnK*, *mtnA*, and *ald2* gene cluster (strain ΔK2), ATCC 25922 was completely incapable of growth with 5dR (Fig. 4A), but could still grow using glucose (Fig. S6). To verify that DHAP shunt inactivation was responsible for the loss of growth using 5dR, we reintroduced the DHAP shunt gene cluster *in trans* from a tetracycline-inducible plasmid, which fully restored aerobic growth with 5dR (Fig. 4A; pK2). This establishes that the DHAP shunt is a carbon assimilation pathway for 5dR as the sole growth substrate under aerobic growth conditions.

**Fig. 4.**
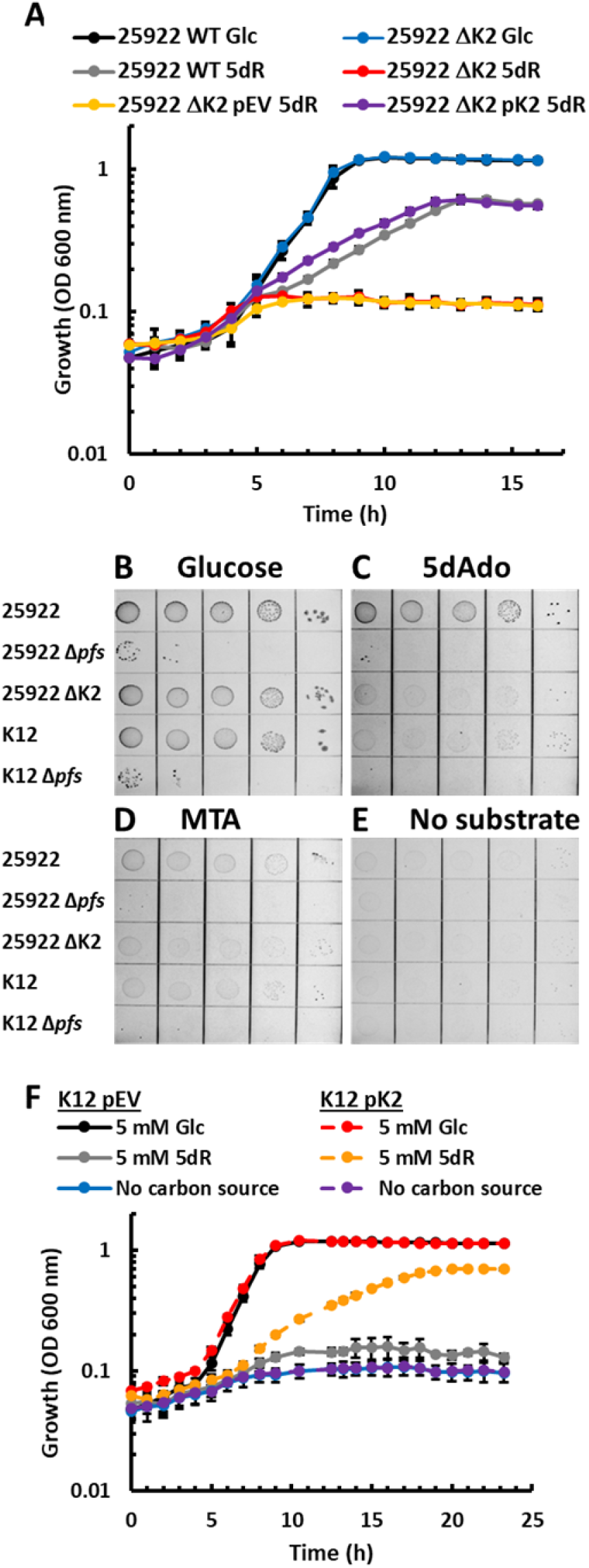
Assimilation of 5’-deoxy-nucleosides and 5-deoxy-pentoses for carbon and energy metabolism by ExPEC with the DHAP shunt. **A)** Growth of either wild type ATCC 25922, ATCC 25922 ΔK2 (Δ*mtnK* Δ*mtnA* Δ*ald2*), ΔK2 + pEV (pTETTET empty vector), or ΔK2 + pK2 (pTETTET vector with DHAP shunt *mtnK, mtnA,* and *ald2*) on either 25 mM glucose (Glc) or 5 mM 5-deoxy-D-ribose (5dR) as the carbon source. **B-E)** Growth of ATCC 25922, ATCC 25922 Δ*pfs*, ATCC 25922 ΔK2, K12 commensal strain BW25113, or BW25113 Δ*pfs* on 1 mM of (B) glucose, (C) 5dAdo, (D) MTA, (E) or no carbon source. **F)** Growth of K12 commensal strain BW25113 pEV (pTETTET empty vector) and K12 complemented with pK2 (pTETTET with DHAP shunt) using either 5 mM glucose (Glc) or 5 mM 5-deoxy-D-ribose (5dR) as the carbon source, or no carbon source. Average and standard deviation error bars are for n=3 independent replicates.

To further quantify the role of the DHAP shunt as a carbon assimilation pathway from 5’-deoxy-nucleosides, we grew ATCC 25922 and K12 with MTA and 5dAdo as a sole carbon source. Given that concentrations of these substrates above 1 mM are inhibitory (Fig. 3), we employed serial dilution plating using M9 agar plates supplemented with 1 mM glucose, MTA, or 5dAdo (Fig. 4B-E) instead of liquid cultures which require ∼5 mM substrate to quantify growth. The ATCC 25922 wild type strain could utilize 5dAdo for growth similar to that of glucose. Growth with MTA was also observable, but poor compared to glucose and 5dAdo. Consistent with the 5dR growth studies (Fig. 4A), deletion of the ExPEC DHAP shunt resulted in minimal growth with 5dAdo and MTA, which was identical to the baseline growth of the K12 commensal strain. Deletion of the Pfs nucleosidase in both strains resulted in poor growth on 1 mM glucose as well as 5dAdo and MTA, underpinning the importance of the nucleosidase in preventing the inhibitory accumulation of MTA and 5dAdo via intracellular SAM-dependent processes (Fig. 3) [17, 38]. Thus, the DHAP shunt enables the metabolism of a diverse number of 5’-deoxy-nucleosides and 5-deoxy-pentoses for use as growth substrates under aerobic growth conditions.

Given that the efficiency by which an organism can convert a growth substrate into cell mass is dependent upon the nature of the growth substrate and how it can be metabolized, we assessed how efficiently *E. coli* could use 5dR versus other known aerobic growth substrates by measuring growth yield, Y_X/S_ (g cell biomass per g substrate) (Fig. 5A). As expected, glycolytic and pentose phosphate substrates resulted in 0.5 g cells per g substrate, and pyruvic acid, which is the end product of glycolysis before entry into the TCA cycle, resulted in a yield of 0.3 g cells per g substrate [44, 45]. The decreased efficiency for compounds of lower glycolysis (pyruvate, lactate, DHAP) is due in part to an increased deleterious flux through the TCA cycle as wasted CO_2_ [44] and/or due to being more oxidized than glucose, requiring a larger percentage of the substrate to be oxidized to CO_2_ by the TCA cycle to generate enough reducing equivalents [NAD(P)H] to build cell biomass [46]. The 5-deoxy-pentose sugar, 5dR, resulted in a yield of 0.3 g per g 5dR, just as for pyruvate. While growth with fucose and rhamnose 6-deoxy-hexoses was more efficient (0.4 g per g substrate), neither 5-deoxy-pentoses nor 6-deoxy-hexoses could be utilized as efficiently as glucose. Thus, as for 6-deoxy-hexoses, the 5-deoxy-pentoses likely serve as secondary sugars when a preferred sugar like glucose is present [47, 48].

**Fig. 5.**
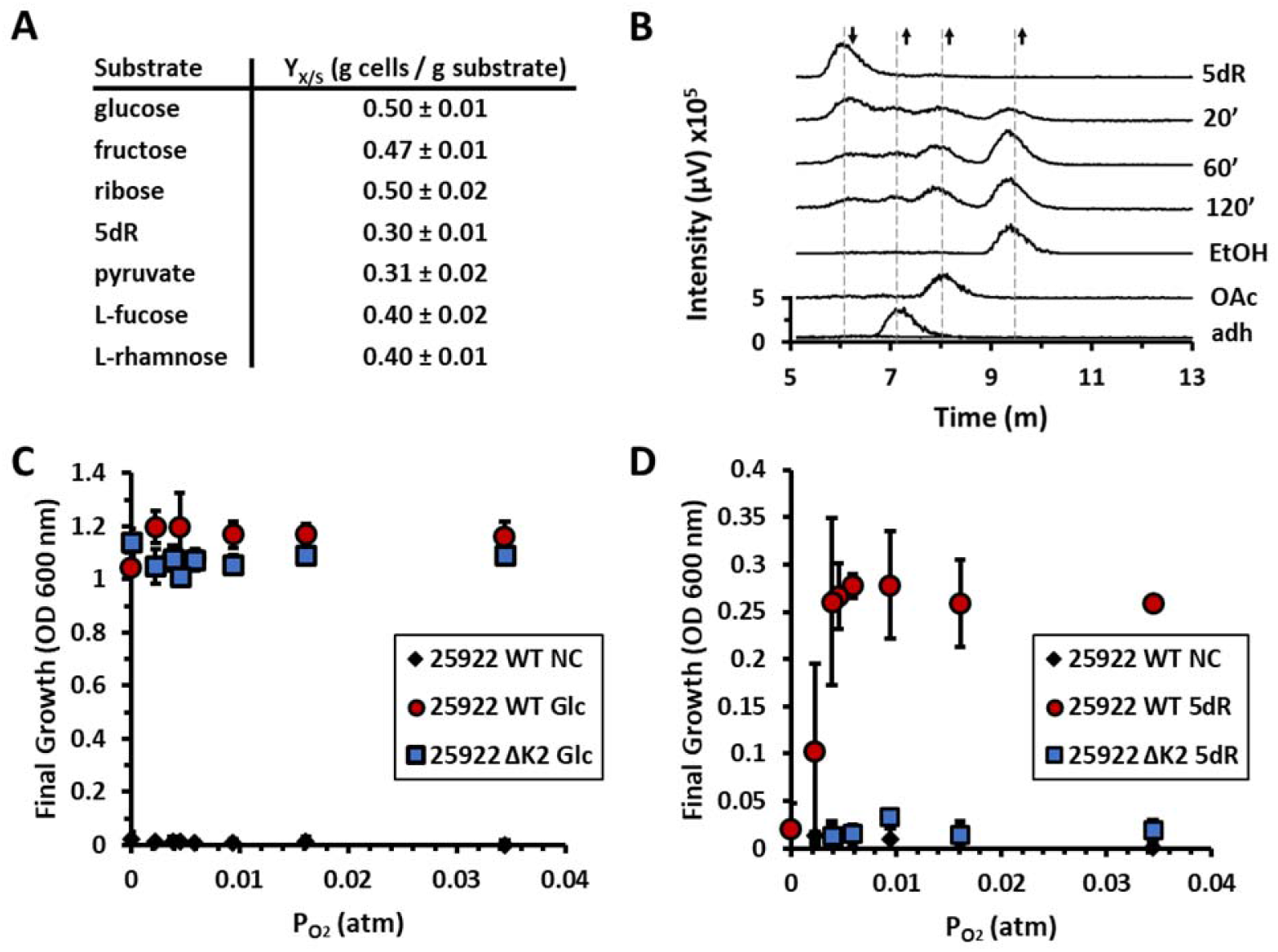
The ExPEC DHAP shunt is an efficient carbon assimilation pathway for growth, but only under aerobic conditions. **A)** Aerobic ATCC 25922 growth efficiency for various carbon growth substrates measured as dry cell weight of cells generated per gram of substrate. **B)** Ion exclusion HPLC quantification of ^3^H-labeled metabolites produced upon feeding *E. coli* ATCC 25922 with [^3^H-methyl]-5’-deoxyadenosine under anaerobic conditions. After feeding, metabolism was quenched by rapid freezing in liquid nitrogen at the indicated time in minutes. **C-D)** ExPEC ATCC 25922 wild type and DHAP shunt deletion (ΔK2) strains were grown under varying oxygen concentrations with either C) 25 mM glucose (Glc) or D) 5 mM 5-deoxy-D-ribose (5dR) as the sole carbon source. In addition, experiments were performed with no carbon source (NC). Final optical density measurements at 600 nm (OD_600_) were taken after 24 hours of growth. Averages and standard deviation error bars in A,C-D are for n=3 independent replicates.

### O_2_ requirements of DHAP shunt in E. coli

ExPEC strains can inhabit a number of niches, including the intestine, urinary tract, and blood, which widely differ in oxygen and nutrient availability. While the intestinal tract is primarily anoxic, urine in healthy individuals typically has a dissolved O_2_ concentration of about 4.2 ppm, which corresponds to ∼0.065 atm (50 mm Hg) oxygen partial pressure [49]. Oxygen concentration can vary with corresponding infection and urine microbial composition [50, 51]. Given that the DHAP shunt supported growth with SAM utilization byproducts as carbon substrates under rigorously aerobic but not under anaerobic conditions, this raised the question as to whether the DHAP shunt could support growth under oxygen tensions similar to those found in urine. Oxygen concentrations were varied from anaerobic to microaerobic to aerobic and cultures were monitored for growth using glucose or 5dR as the sole carbon source (Fig. 5C-D). The ATCC 25922 wild type and DHAP shunt deletion strain (strain ΔK2) were able to grow normally using glucose across all oxygen concentrations tested (Fig. 5C). In stark contrast, ATCC 25922 wild type strain could only grow using 5dR as a sole carbon source at oxygen tensions ≥ 0.004 atm (3 mm Hg). Addition of nitrate for anaerobic respiration did not allow for anaerobic growth using 5dR. The observed growth with 5dR at the various oxygen tensions required the DHAP shunt, as no growth was observed with 5dR in the DHAP shunt deletion strain (Fig. 5D). We verified that the DHAP shunt was active under anaerobic growth conditions through targeted metabolomics assays. Cells grown on glucose anaerobically and then fed with ^3^H-labeled 5dAdo showed conversion to 5dR and ultimately a mixture of acetate, acetaldehyde, and ethanol (Fig. 5B). This contrasts with aerobic growth in which only ethanol is formed [13]. Ultimately, the DHAP shunt appears to be physiologically relevant across the range of oxygen tensions found in typical extraintestinal environments for use of SAM utilization byproducts as growth substrates. Conversely, under anaerobic conditions *E. coli* does not appear to be able to perform sufficient fermentation or anaerobic respiration with these compounds via the DHAP shunt for growth.

### Introduction of the DHAP shunt to commensal E. coli enables growth with SAM utilization byproducts

The genes encoding *mtnK*, *mtnA*, and *ald2* of the DHAP shunt are not present in commensal *E. coli* and as such, commensal *E. coli* cannot grow using MTA, 5dAdo, or 5dR as the sole carbon source (Fig. 4). Given the ExPEC DHAP shunt is located on the tRNA-*leuX* genomic island [13], we questioned if there are any other factors present in ExPEC strains that are not present in commensals which are required for growth using SAM utilization byproducts. We introduced the DHAP shunt gene cluster into *E. coli* K12 and expressed the genes *in trans* from the same tetracycline-inducible plasmid used in ATCC 25922. The expression of *mtnK*, *mtnA*, and *ald2* enabled aerobic growth of commensal strain K12 with 5dR (Fig. 4F). Thus, for aerobic growth and energy metabolism, *E. coli* can gain the ability to use SAM utilization byproducts through acquisition and expression of the DHAP shunt genes *mtnK*, *mtnA*, and *ald2* without any additional factors.

## Discussion

SAM is a ubiquitous cofactor used by all organisms for methylations, polyamine synthesis, quorum sensing, and radical enzyme chemistry, which results in the formation of SAH, MTA, and 5dAdo (Fig. 1) [52]. Subsequently, all organisms must eliminate these inhibitory compounds from the cell through further metabolism into less inhibitory species and/or excretion [23]. While SAH is universally recycled by all organisms (except for some obligate endosymbionts) by a variation of the active methyl cycle (Fig. 1) [23], recent studies have revealed that a diversity of pathways exist for the detoxification and recycling of MTA (Suppl. Fig. S1) [22, 32, 33, 53–55]. These pathways can be bifunctional for both MTA and 5dAdo as in the case of the DHAP shunt [13, 15], or promiscuous for adenine as in the case of the Universal Methionine Salvage pathway [56]. Historically, metabolism of MTA has been primarily considered to be a detoxification process to prevent inhibitory accumulation, and if recycled to methionine (Suppl. Fig. S1), a sulfur salvage process with tie-ins to formate, isoprenoid, and ethylene synthesis [14, 22, 23, 32, 34]. Analogously, metabolism of 5dAdo has been considered to be a detoxification process [15, 17, 18], with tie-ins to 6-deoxy-5-ketofructose-1-phosphate production for amino acid biosynthesis in *M. jannaschii* and 7-deoxysedoheptulose production as an anti-microbial secondary metabolite in *Synechococcus sp.* [57–59].

Unlike organisms in which the DHAP shunt was initially discovered to function as a methionine salvage pathway via formation of 2-methylthioethanol I as an intermediate (Fig. S1) [13, 14, 20–22], *E. coli* does not appear to use the DHAP shunt for sulfur salvage and methionine synthesis from MTA (Fig. 2, Suppl. Fig. S3D). Furthermore, in clear contrast to the insect pathogen, *Bacillus thuringiensis* [15], the DHAP shunt kinase (MtnK), isomerase (MtnA), and aldolase (Ald2) do not appear essential for the elimination of inhibitory byproducts of SAM utilization (Fig. 3). These DHAP shunt genes unique to ExPEC strains and their deletion had no effect on the sensitivity of *E. coli* to MTA, 5dAdo, and 5dR. Only the Pfs nucleosidase, which is conserved across all *E. coli* for the hydrolysis of SAH, MTA, and 5dAdo, was absolutely required to prevent growth inhibition in the presence of these compounds. Thus, the conserved Pfs nucleosidase is the primary means for managing inhibitory buildup of SAH, MTA, and 5dAdo produced by SAM-dependent enzymes. Coordinately, MTR and 5dR do not appear to need to be managed beyond export from the cell [16, 17].

The prime function of the DHAP shunt in *E. coli* is evidently for carbon assimilation and energy metabolism from 5’-deoxy-nucleoside and 5-deoxy-pentose sugars (Fig. 4). The pathway does have a preference for 5dAdo and 5dR versus MTA and MTR as growth substrates, which is corroborated by our previous enzymatic analyses showing that the *E. coli* ATCC 25922 kinase (MtnK) had a 10-fold higher specificity for 5dR versus MTR (k_kat_/K_M_ = 6.3×10^5^ vs. 0.6×10^5^ M^-1^s^-1^) [13]. Under aerobic conditions the acetaldehyde produced from 5’-deoxy-nucleoside and 5-deoxy-pentose substrates accumulates as ethanol as previously shown [13]. This is because during aerobic growth, *E. coli* cannot use acetaldehyde and ethanol as growth substrates due to oxygen inactivation of the trifunctional CoA-acylating AdhE aldehyde/alcohol dehydrogenase for metabolism of ethanol and acetaldehyde to acetyl-COA. Rather, the acetaldehyde can only be converted to ethanol as a terminal product by alcohol dehydrogenase AdhA [60, 61]. Interestingly the DHAP shunt is active under anaerobic conditions, as can be seen from the metabolism of 5dR into ethanol, acetaldehyde, and acetate (Fig. 5B), but evidently is unable to support growth for reasons currently unknown. The mixture of acetaldehyde, ethanol and acetate is due to AdhE activity, which reduces acetaldehyde to ethanol or oxidizes it to acetyl-CoA, depending on NAD+/NADH availability. Subsequently, the acetyl-CoA can be further metabolized to acetate by the phosphotransferase pathway [60, 61].

Strikingly, the inability of the DHAP shunt to serve as an anaerobic fermentative or respiratory carbon assimilation and energy production pathway in *E. coli* sheds light on the disparity as to why the DHAP shunt is found exclusively in ExPEC strains and not in intestinal commensal and pathogenic *E. coli* [13]. While the urine, blood, and mammary niches where *ExPEC* strains can inhabit are oxygenated [49, 62], the large intestine is predominantly anoxic. Even during infection, which can cause O_2_ levels to drop in urine, blood, and milk [50, 51, 62], the resulting lower oxygen tensions are still sufficient to enable functionality of the DHAP shunt as a carbon assimilation and energy metabolism pathway (Fig. 5). Thus, it appears that the DHAP shunt likely would not confer any significant growth advantage in the intestinal environment even though 5’-deoxy-nucleosides or 5-deoxy-pentose sugars are certainly present. Strikingly, in our previous comparative genomic analyses [13], the DHAP shunt was invariably located in ExPEC strains at the distal end of the tRNA-*leuX* genomic island adjacent to the *yjhUFGHI* and *sgcREAQCBX* gene clusters for putative xylonate metabolism and PTS sugar transport that is conserved across all *E. coli*. Given that ExPEC strains are viewed to have evolved from commensals through horizontal gene acquisition of elements that confer pathogenicity or fitness in the extraintestinal niche [63], it appears that *E. coli* may have acquired the DHAP shunt early on during its transition to the extraintestinal lifestyle. Also, from this point of view, the lack of DHAP shunt utility for *E. coli* in anoxic environments points to why intestinal *E. coli* have not acquired the pathway. Given that the DHAP shunt is widespread across bacteria and particularly enriched in pathogenic species of many genera, including *Bacillus* and *Clostridium sp.* (e.g., *B. cereus*, *B. anthracis*, *C. botulinum*, *C. tetani*), these findings that the DHAP shunt can function as a carbon assimilation and energy metabolism pathway call for deeper investigation in both aerobic and anaerobic pathogens on the role of the DHAP shunt beyond serving as a 5’-deoxy-nucleoside and 5-deoxy-pentose detoxification pathway as initially proposed [15].

## Acknowledgments

The authors thank the OSU Campus Chemical Instrument Center and Dr. Wu Gong for providing untargeted metabolomics support and use of the Thermo QE M UPLC MS/MS System. These resources are supported in part by the NIH Award Number P30 CA016058. This work was funded by the NIH NIAID grant 1R01AI154456-01 (J.A.N. and F.R.T).

## Author Contributions

K.A.H., J.T.G., J.A.W., and J.A.N. performed the growth and genetic analyses. K.A.H. performed the untargeted and targeted metabolomics and J.T.G. performed the metabolite quantification analyses. J.A.N. and F.R.T designed the experiments and acquired funding. J.A.N. supervised the research. All authors discussed the research and contributed to writing the manuscript.

## Supplementary Information Supplementary Methods

### Construction of pTETTET complementation vector

Plasmid pTETTET, with a tetracycline-inducible promoter, was constructed by replacing the *lac* promoter of pBBRsm2-MCS6 [29] with the dual *tet* promoter plus *tetR* and the 70 bp region immediately downstream of *tetR* from the transposon Tn*10* [30, 31]. The insertion sequence was synthesized (Genewiz, Azenta Life Sciences), which included flanking NdeI and AseI restriction sites for cloning into pBBRsm2-MCS6. In the synthesized fragment (see below sequence), internal NdeI and AseI sites in the *tet* regulon were removed such that only silent mutations were introduced into the *tetR* sequence. The *tet* regulon was amplified using primers Pzt1RBS-AseI-F and Pzt1RBS-NdeI-R (Suppl. Table S1) and inserted into pBBRsm2-MCS6 to form pTETTET such that the *tetA* promoter was upstream of *lacZa* and the MCS of pBBRsm2-MCS6 (Suppl. Fig. S2).

### Synthesized Tet Promoter Sequence (with AseI and NdeI sites underlined)

#### attaat

TCTAAAGGGTGGTTAACTCGACATCTTGGTTACCGTGAAGTTACCATCACGGAAAAAGGTTATGCTGCTTTTAAGACCCACTTTCACATTTAAGTTGTTTTTCTAATCCGCAGATGATCAATTCAAGGCCGAATAAGAAGGCTGGCTCTGCACCTTGGTGATCAAATAATTCGATAGCTTGTCGTAATAATGGCGGCATACTATCAGTAGTAGGTGTTTCCCTTTCTTCTTTAGCGACTTGATGCTCTTGATCTTCCAATACGCAACCTAAAGTAAAATGCCCCACAGCGCTGAGTGCATATAATGCATTCTCTAGTGAAAAACCTTGTTGGCATAAAAAGGCTAATTGATTTTCGAGAGTTTCATACTGTTTTTCTGTAGGCCGTGTACCTAAATGTACTTTTGCTCCATCGCGATGACTTAGTAAAGCACATCTAAAACTTTTAGCGTTATTACGTAAAAAATCTTGCCAGCTTTCCCCTTCTAAAGGGCAAAAGTGAGTATGGTGCCTATCTAACATCTCAATGGCTAAGGCGTCGAGCAAAGCCCGCTTATTTTTTACATGCCAATACAATGTAGGCTGCTCTACACCTAGCTTCTGGGCGAGTTTACGGGTTGTTAAACCTTCGATTCCGACCTCATTAAGCAGCTCTAATGCGCTGTTAATCACTTTACTTTTATCTAATCTAGACATCgTTAATTCCTAATTTTTGTTGACACTCTATCATTGATAGAGTTATTTTACCACTCCCTATCAGTGATAGAGAAAAGTc

#### atatg

### Radiometric Targeted Metabolomics

For 14C-labeled MTA feedings, cells were grown aerobically to mid-log phase in M9 glucose media, washed 3 times with sulfur-free M9 glucose media, and resuspended to an OD_600nm_ ∼5 in sulfur-free M9 glucose media. Cells were fed with a mixture of 15 μM [^14^C-methyl]-5’-methylthioadenosine (0.1 μCi/100 μl) and 200 μM MTA, and 100 μl aliquots were placed in an open 2 ml conical glass tube, bubbled with purified air, and incubated at 120 RPM at 37 °C in a shaking water bath. Samples (100 μl) were collected at indicated timepoints and flash frozen in LN_2_. Metabolites were collected by thawing the cells to room temperature, vortexing, centrifuging at 5000 x g for 1 minute, and collecting the spent media supernatant. Metabolites were resolved by C18 reverse phase HPLC with inline liquid scintillation detection [13]. [^14^C-methyl]-5’-methylthioadenosine was synthesized from [^14^C-methyl]-*S*-adenosyl-L-methionine (Perkin-Elmer) by acid hydrolysis [13]. Similarly, for 5dAdo feedings, anaerobically grown cells were washed with carbon-free M9 media and resuspended to an OD_600nm_ ∼5 in carbon-free M9 media. Cells were fed with a mixture of 0.4 μM [5’-^3^H]-5’-deoxyadenosine (0.2 μCi/100 μl) and 200 μM 5dAdo and incubated in the shaking water bath as 100 μl aliquots in 2 ml sealed serum vials. Cells were collected at indicated timepoints by centrifugation at 5000 x g for 1 minute, and metabolites present in the spent media were resolved by ion exclusion chromatography with inline scintillation detection [13].

## Supplementary Results

### Recycling of endogenous SAM utilization byproducts is not a significant role of the DHAP shunt in ExPEC

In addition to using the DHAP shunt for assimilation of exogenous SAM utilization byproducts from the environment, ExPEC strains could use the DHAP shunt to salvage endogenously produced SAM utilization byproducts to support growth, particularly under carbon-or sulfur-limiting conditions. We determined if ExPEC strain ATCC 25922 produced more MTA and 5dAdo than its commensal counterpart, K12, by inactivating the Pfs nucleosidase, which prevents further metabolism of MTA, 5dAdo, and SAH. Subsequent accumulation and quantification of these compounds in the surrounding media enables relative comparison of SAM byproducts produced (Suppl. Fig. S3A-B) [13]. During aerobic growth, as evidenced by the Pfs deletion, ATCC 25922 produced twofold more MTA than commensal strain K12 and twofold less SAH (Suppl. Fig. 3A; t-test, P<0.05). The low amounts of 5dAdo produced aerobically are due to the general oxygen sensitivity of radical SAM enzymes which limits their use aerobically. For anaerobic growth there was no statistically significant difference in the amounts of SAH, MTA, and 5dAdo produced between ATCC 25922 and K12 (Suppl. Fig. S3B; t-test, P>0.5). Thus, save for a twofold increase in MTA production as observed for other intestinal pathogenic *E. coli* [36], ExPEC strain ATCC 25922 does not appear to produce substantially more SAM utilization byproducts that its commensal counterpart.

To determine if the formation of these SAM utilization byproducts resulted in a metabolic burden on carbon or sulfur pools of the ExPEC strain that was alleviated through salvage, we compared the growth of wild type ATCC 25922 and the DHAP shunt deletion strain (strain ΔK2) under replete and limiting glucose or sulfate conditions (Suppl. Fig. S3C-D). There was no difference in growth rate or final cell density between the ATCC 25922 wild type and DHAP shunt deletion strains for each glucose or sulfate concentration. Taken together with the SAM utilization byproduct quantification (Suppl. Fig. S4A-B), these results indicate that ExPEC strain ATCC 25922 does not produce substantial amounts of endogenous MTA and 5dAdo such that the DHAP shunt is required as a salvage pathway to prevent detrimental carbon or sulfur loss. However, this does not exclude such a role for the DHAP shunt in other ExPEC strains or in a particular infection niche.

## Supplementary Tables

**Supplementary Table S1.**
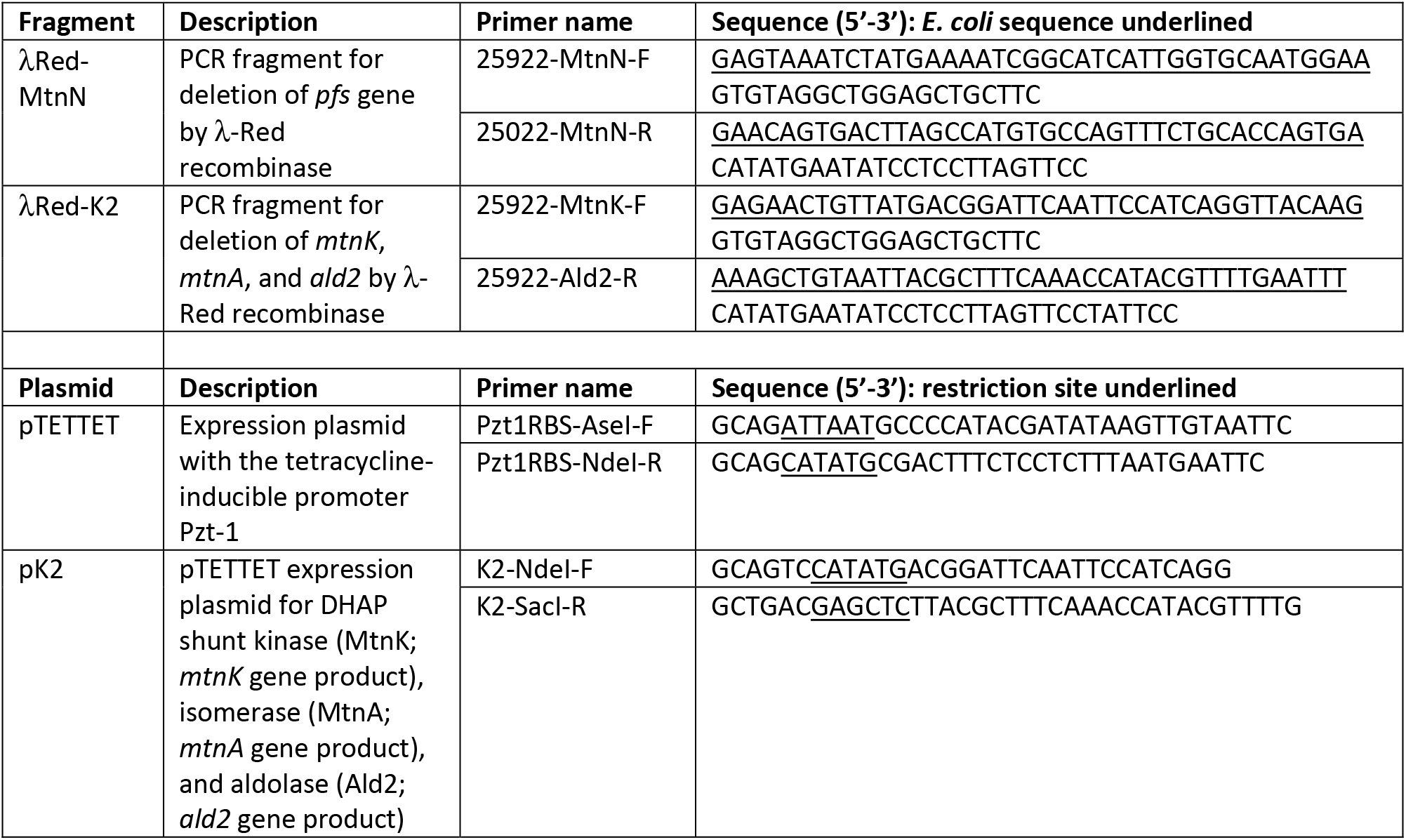
Primers and Plasmids Used in this Study.

## Supplementary Figures

**Suppl. Fig. S1.**
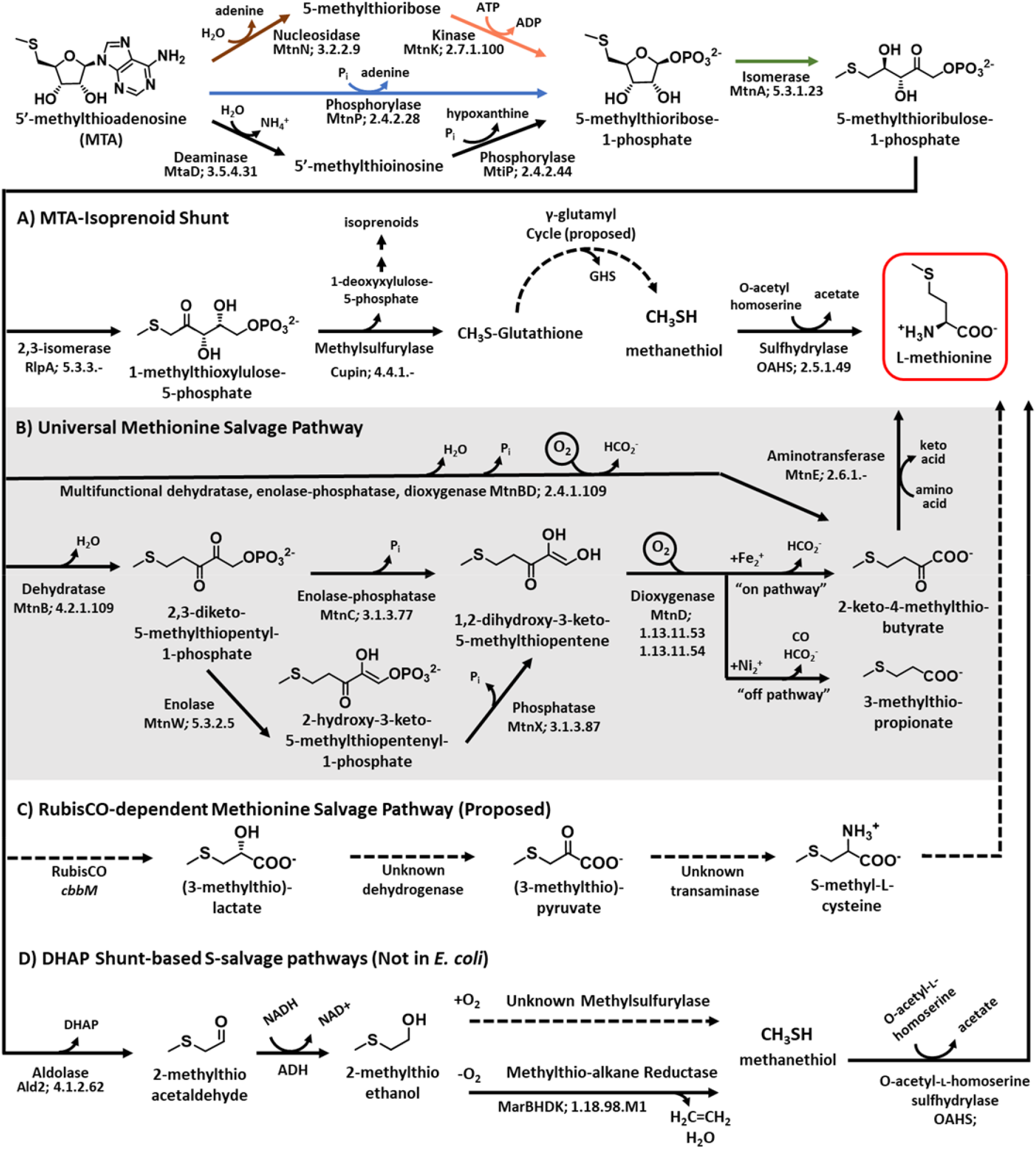
Diversity of sulfur salvage pathways for MTA present in various bacteria. While *E. coli* possesses the DHAP shunt (MtnN, MtnK, MtnA, and Ald2) as well as an alcohol dehydrogenase to ultimately convert MTA to 2-methylthioethanol, it is missing gene homologs for other known MTA salvage pathways other organisms (A-D). All known sulfur salvage pathways from MTA begin by nucleoside cleavage. Some organisms such as *P. aeruginosa* and *M. jannaschii* initially deaminate MTA to enable specific nucleoside cleavage [57, 58, 66, 67]. **A)** The MTA-isoprenoid shunt from *R. rubrum* leads to the formation of isoprenoids and methanethiol as the immediate methionine precursor [21, 32, 37, 68]. **B)** Microbial variations of the Universal Methionine Salvage Pathway leads to formate and 2-keto-4-methylthiobutyric acid as the immediate methionine precursor [34, 53–55]. **C)** The proposed RubisCO-dependent MTA metabolism pathway from *R. rubrum* [33, 69]. **D)** Several sulfur salvage pathways exist that utilize 2-methylthioacetaldehyde produced by the DHAP shunt. In *R. palustris* under aerobic growth conditions (+O_2_), an unknown methylsulfurylase cleaves 2-methylthioethanol into methanethiol as the immediate methionine precursor [20]. Under anaerobic conditions (-O_2_), organisms with the nitrogenase-like methylthio-alkane reductase reductively cleave 2-methylthioethanol into methanethiol and ethylene[14, 22]. Protein designations and EC numbers are provided below each enzymatic step. ADH, unknown alcohol dehydrogenase.

**Suppl. Fig. S2.**
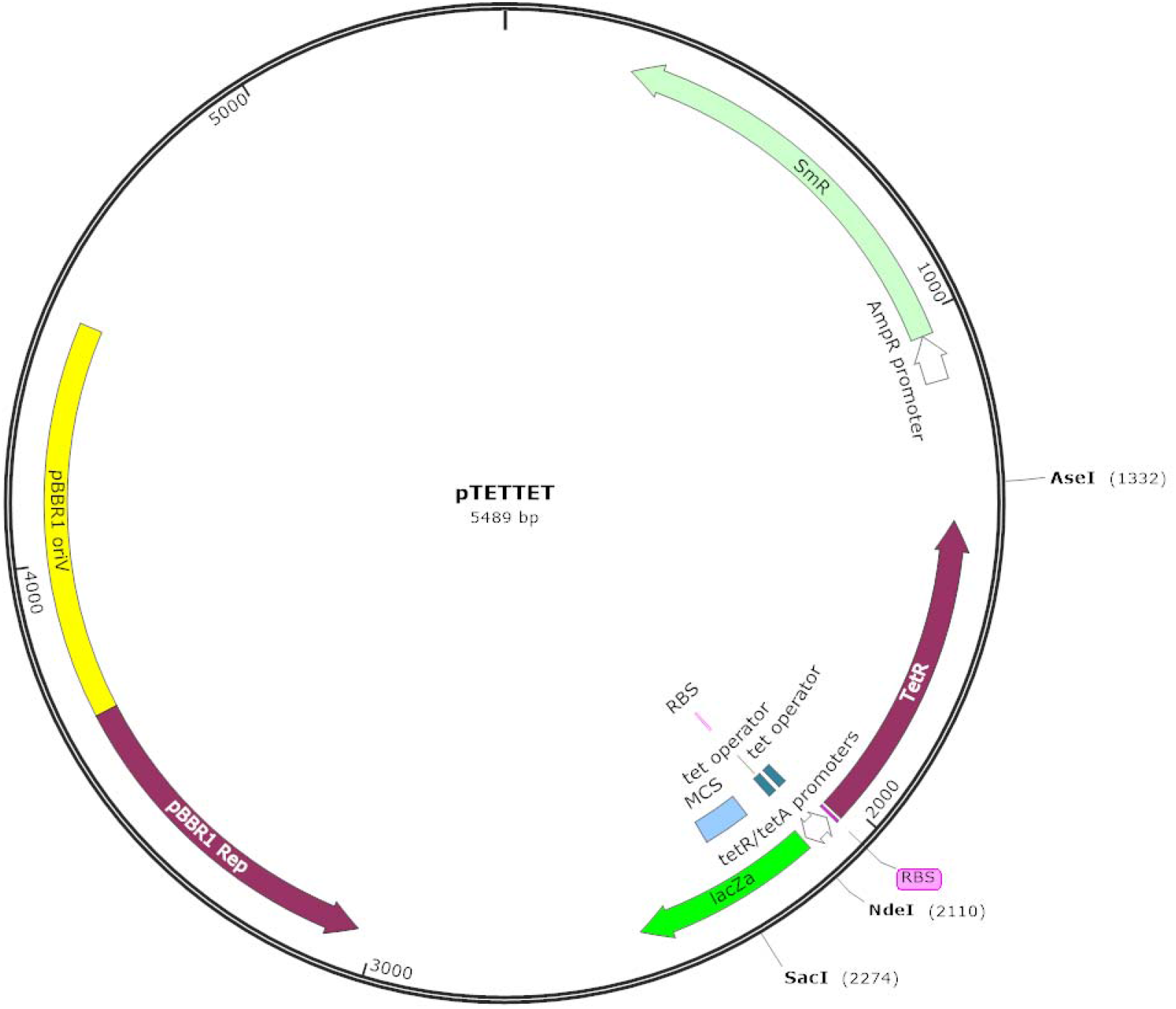
pTETTET plasmid map for complementation in *E. coli*. The dual Tet promoter and *tetR* sequence from the transposon Tn10 [30, 31] was synthesized and modified to contain flanking NdeI and AseI sites. Internal NdeI and AseI sites from TetR were removed by silent substitutions (sequence in Suppl. Methods). This modified Tet regulon was amplified with primers Pzt1RBS-AseI-F and Pzt1RBS-NdeI-R (Suppl. Table S1) and inserted between the NdeI and AseI sites of pBBRsm2-MCS6 to form plasmid pTETTET. Plasmid map constructed with SnapGene.

**Suppl. Fig. S3.**
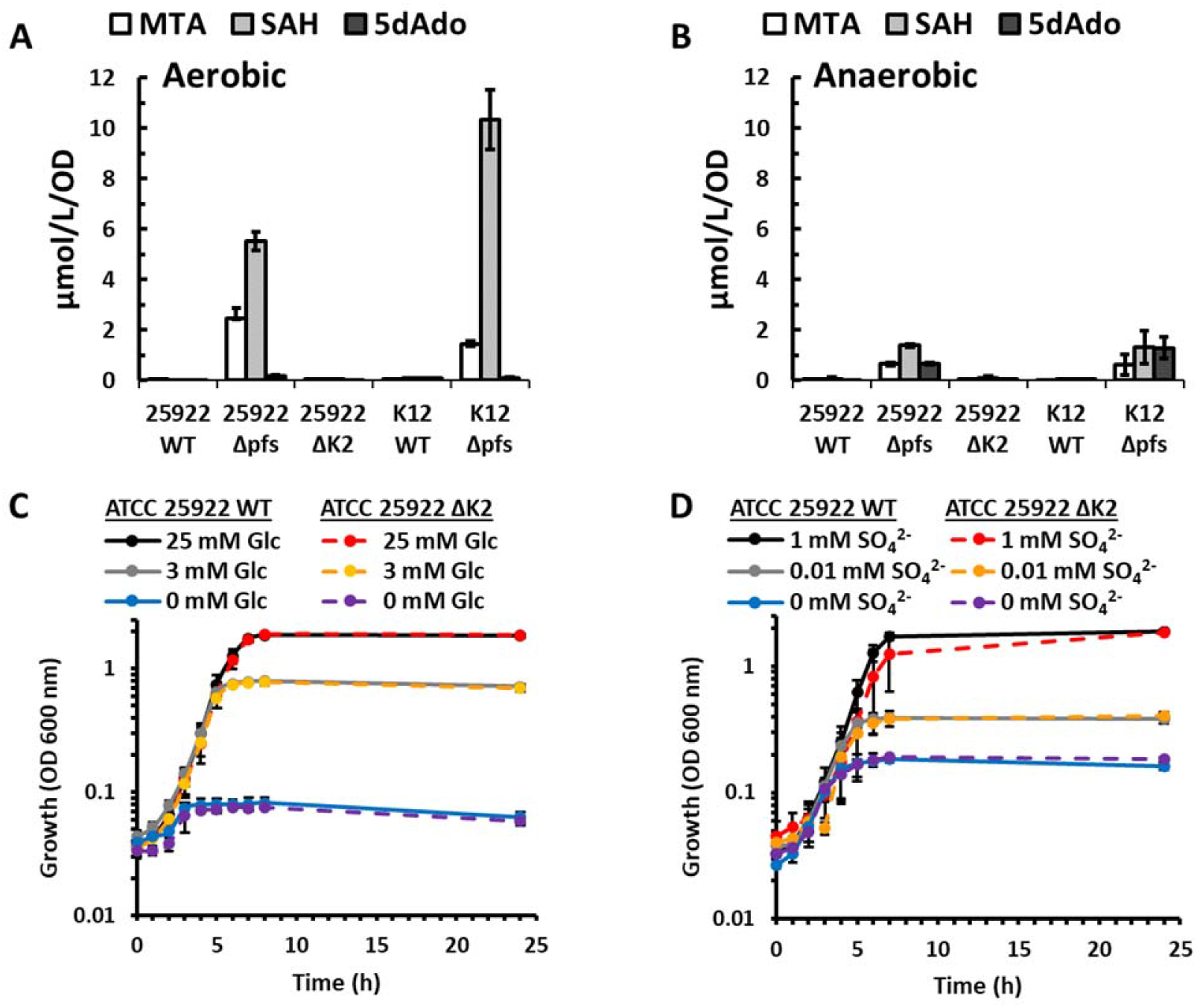
Intracellular SAM byproducts do not need to be salvaged by the DHAP shunt to maintain *E. coli* growth. A-B) HPLC analysis of internal DHAP shunt metabolites from cultures grown with 25 mM glucose as the sole carbon source under (A) aerobic or (B) anaerobic conditions. *E. coli* ATCC 25922 or K12 cultures were harvested at mid-log phase for metabolite analysis. WT, wild type; Δ*pfs*, Pfs nucleosidase deletion strain; ΔK2, *mntK*-*mtnA*-*ald2* deletion strain. **C-D)** Growth of either wild type ATCC 25922 or ATCC 25922 ΔK2 strain under decreasing concentrations of (C) glucose or (D) sulfate. Average and standard deviation error bars are for n=3 independent replicates.

**Suppl. Fig. S4.**
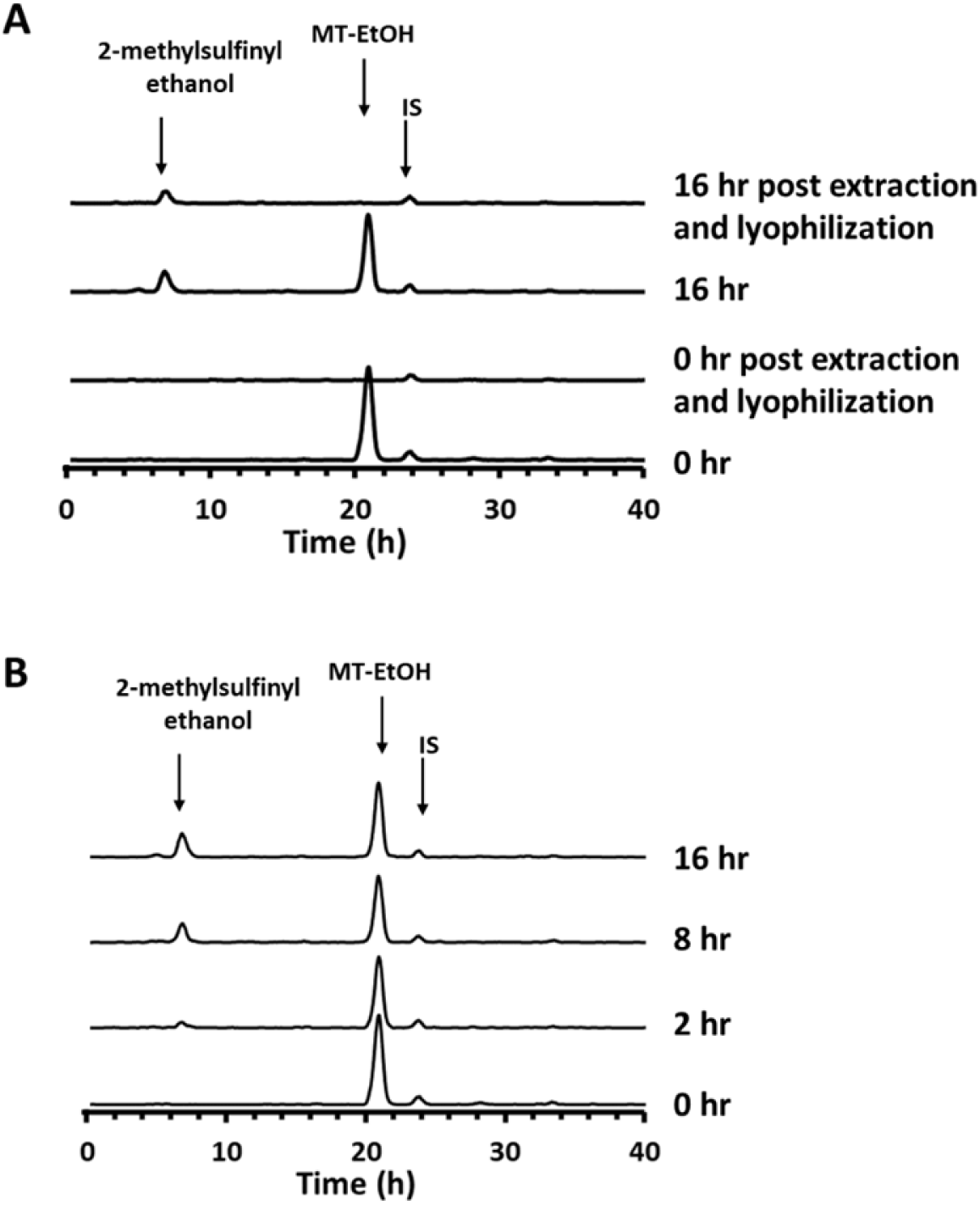
Reverse Phase HPLC quantification of non-enzymatic oxidation of 2-methylthioethanol to 2-methylsulfinylethanol and selective removal of 2-methylthioethanol by lyophilization. **A)** [^14^C-methyl]2-methylthioethanol was synthesized from [^14^C-methyl]-5’-methylthioadenosine as previously described [22] and then incubated in M9 glucose media at 37 °C under aerobic conditions for the indicated amount of time (0 and 16 h) in duplicate. After incubation, one of the two aliquots was combined with an equal volume of acetonitrile as done for extraction of metabolites from cells for LC-MS/MS, centrifuged, and the supernatant lyophilized. Metabolites were resolved by reverse phase HPLC as in Suppl. Fig. S3. **B)** To verify that the oxidation of 2-methylthioethanol to 2-methylsulfinylethanol was temporal, aliquots of [^14^C-methyl]2-methylthioethanol were incubated as in (A) and then resolved by reverse phase HPLC. 2-methylsulfinylethanol, R_T_ = 6.7 min; MT-EtOH – 2-methylthioethanol, R_T_ = 21.7 min; IS - internal standard, R_T_ = 24.0 min.

**Suppl. Fig. S5.**
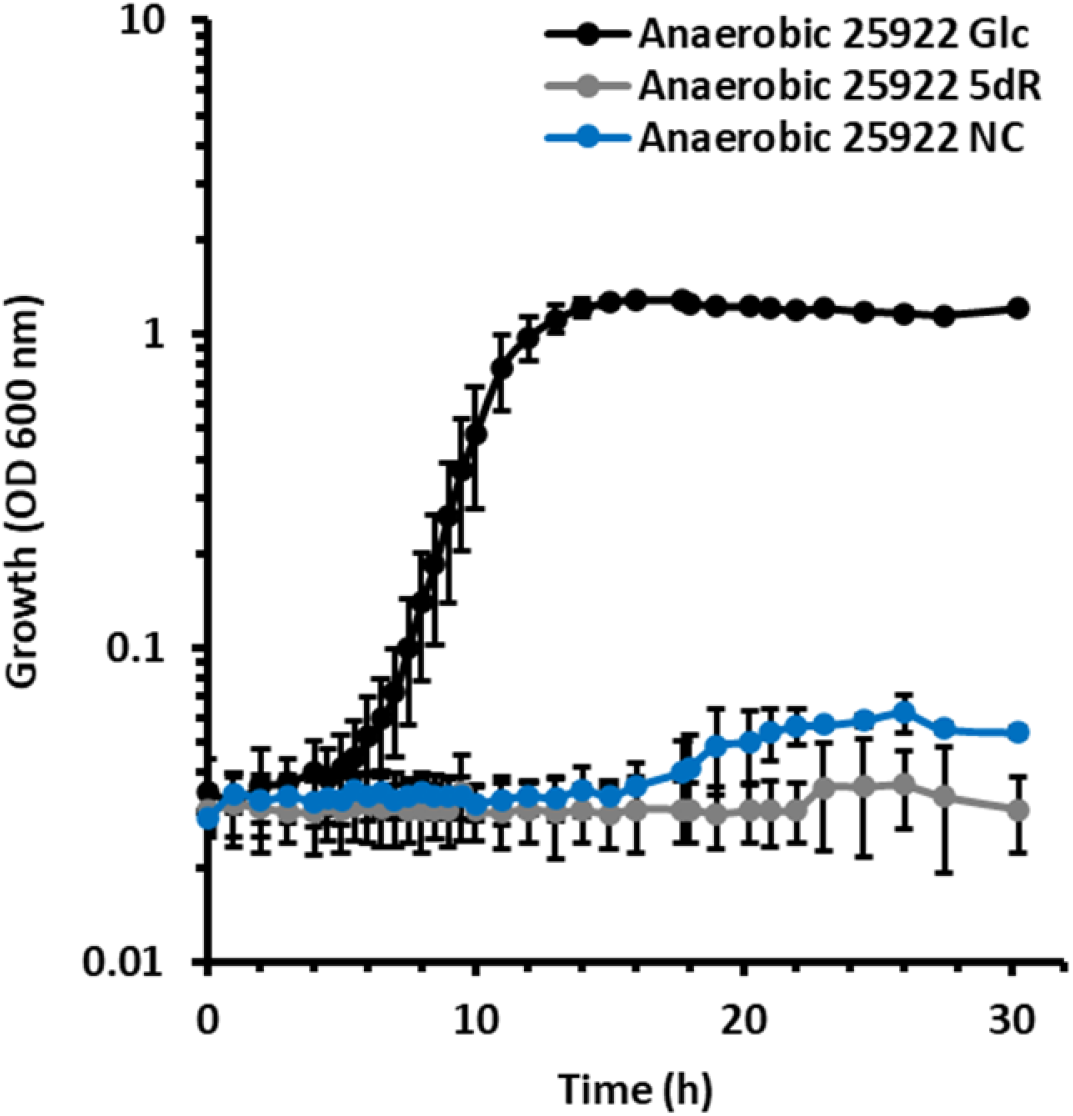
ExPEC ATCC 25922 cannot grow anaerobically with 5dR as a sole carbon source. ATCC 25922 wild type strain was grown anaerobically with no carbon (NC), 5 mM glucose (Glc) or 5 mM 5-deoxy-D-ribose (5dR). Averages and standard deviation error bars are for n=3 independent growth experiments.

**Suppl. Fig. S6.**
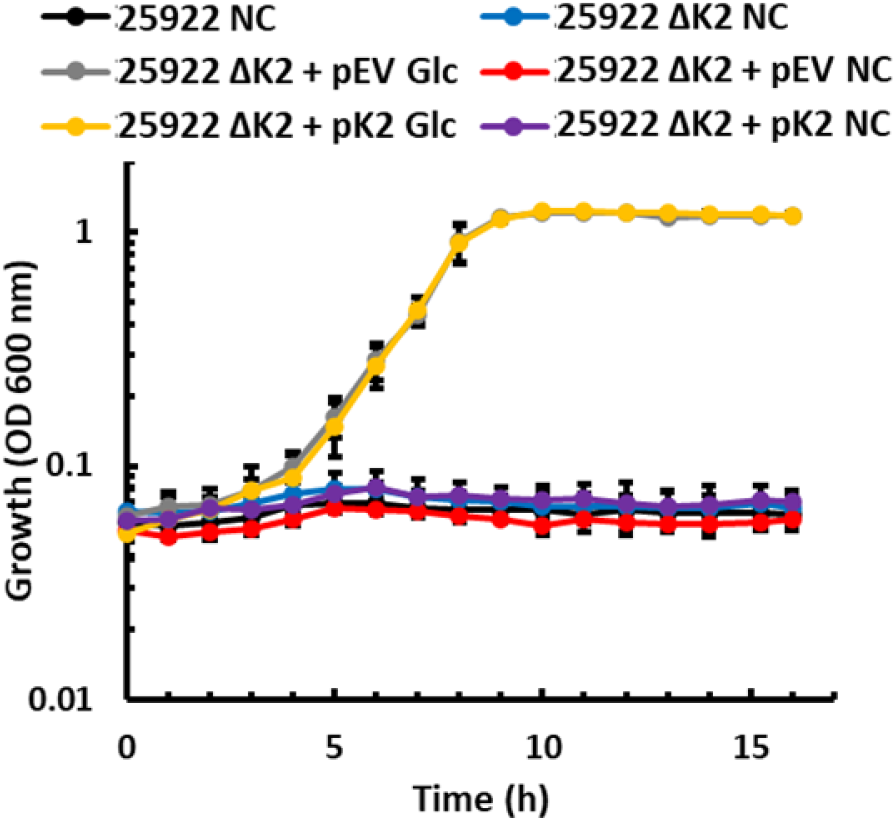
Deletion of DHAP shunt in ExPEC ATCC 25922 does not affect growth on glucose. ATCC 25922 wild type strain and ATCC 25922 DHAP shunt deletion strain (ΔK2) were grown in no carbon media (NC) or in the presence of glucose (Glc). Strains were complemented with either the pTETTET empty vector (pEV) or pTETTET expressing the DHAP shunt genes (pK2). Averages and standard deviation error bars are for n=3 independent growth experiments.

